# Live high-content imaging with automated analysis reveals mitochondrial changes during vascular calcification

**DOI:** 10.64898/2026.02.03.703459

**Authors:** Ji-Eun Lee, Agustina Salis Torres, Abhinanda R. Punnakkal, Xiao Tan, Justyna Cholewa-Waclaw, Amirhossein Kardoost, Qiyu Tang, Craig Leighton, Janelle Leong, Daniel B. Greenslade, Dilip K. Prasad, Robert K Semple, Alison N. Hulme, Carsten Marr, Mathew H. Horrocks, Vicky E. MacRae

**Affiliations:** The Roslin Institute and R(D)SVS, University of Edinburgh, Easter Bush, Midlothian EH25 9RH, UK; IRR Chemistry Hub, Institute for Regeneration and Repair, University of Edinburgh, Edinburgh EH16 4UU, UK; EaStCHEM School of Chemistry, University of Edinburgh, Edinburgh EH9 3FJ, UK; Department of Computer Science, UiT-The Arctic University of Norway, Tromsø, Norway; Computational Health Center, Helmholtz Zentrum München– German Research Center for Environmental Health, Ingolstädter Landstr. 1, 85764 Neuherberg, Germany; Centre for Cardiovascular Science, Queen’s Medical Research Institute (QMRI), The University of Edinburgh, 47 Little France Crescent, Edinburgh EH16 4TJ, UK; School of Life Sciences, Faculty of Science and Engineering, Anglia Ruskin University, Cambridge, CB1 1PT, UK

## Abstract

Mitochondrial dysfunction is implicated in a wide range of disorders, including cancer, neurodegeneration, and cardiovascular diseases. Conventional assays typically assess mitochondrial function by measuring bulk respiration rates across thousands of cells. While informative, these approaches cannot resolve the behavior of individual mitochondria, may overlook rare mitochondrial events, and often lack sensitivity to subtle changes. Fluorescence microscopy provides single-organelle resolution but is usually low throughput; for example, imaging 1000 cells can require from 20 minutes to an entire day depending on imaging mode. Additionally, analysis of fluorescence images frequently relies on manually selected thresholds, introducing potential bias and variability due to differences in signal-to-background ratios across cells. Here, we present a High-throughput Mitochondrial Imaging Platform (H-MIP) combined with a deep learning segmentation model, enabling rapid imaging and automated characterization of mitochondrial morphology in millions of mitochondria from tens of thousands of cells. We demonstrate the utility of H-MIP by assessing mitochondrial morphology following treatment with Mdivi-1, an inhibitor of mitochondrial division. Furthermore, we employ an *in vitro* disease model (vascular calcification) to show that mitochondria become elongated in vascular smooth muscle cells undergoing pathological calcification. Overall, H-MIP provides a scalable and robust tool for investigating mitochondrial structure, with the potential to accelerate therapeutic discovery targeting mitochondrial health.

**Highlights:**

- H-MIP enables high-throughput imaging of mitochondrial morphology and automated single-mitochondrion analysis at scale.
- H-MIP links mitochondrial morphology to an established disease model, with exciting implications for drug discovery.

## Introduction

In addition to their important role as the powerhouse of cells, mitochondria are now recognized as central regulators of cellular metabolism, oxidative stress, calcification processes, and apoptosis (Chen et al., 2023; Kowalczyk et al., 2021; Ott et al., 2007; Wang et al., 2020, Phadwal et al., 2021). Their highly dynamic network architecture, maintained through a balance of fission and fusion, is tightly linked to mitochondrial function (Al Ojaimi et al., 2022). Disruption of the normally tightly regulated network dynamics contributes to mitochondrial dysfunction, a hallmark of numerous common pathologies, including Parkinson’s disease (PD), cancer, chronic kidney disease (CKD), diabetes, and vascular calcification (VC) (Choi et al., 2022, Galvan et al., 2017; Hattori & Mizuno, 2002; Hsu et al., 2016; Iheagwam et al., 2025; Salis Torres et al., 2024; Zeng et al., 2022; Zong et al., 2024). Despite these well-established links, there remains a critical need for robust, quantitative methods to interrogate how changes in mitochondrial morphology drive disease mechanisms and open avenues for therapeutic intervention. Traditionally, mitochondrial function has been assessed using bulk bioenergetic assays that measure ATP production, oxygen consumption rate, and extracellular acidification rate (Desousa et al., 2023). While these methods provide valuable insights into metabolic activity and energy-generating capacity, they lack single-organelle resolution, and neglect the structural diversity of individual mitochondria. They also only assess mitochondria as only energy-generating organelles, neglecting their other important roles. As a result, early and disease-relevant changes in mitochondrial morphology, often reflecting functional decline, may go undetected.

Fluorescence microscopy overcomes these limitations by allowing visualization of mitochondrial architecture after labeling with dyes such as tetramethylrhodamine methyl ester (TMRM) and MitoTracker (Desai et al., 2024; Neikirk et al., 2023). Unlike bulk assays, fluorescence microscopy reveals detailed information about mitochondrial shape, size, and network organization, features that are increasingly recognized as critical determinants of cellular health and disease (Rafelski, 2013; Kritskaya et al., 2024). Additionally, fluorescence microscopy enables the study of dynamic processes such as mitochondrial fission and fusion (Song et al., 2008). There are, however, some limitations in using fluorescence microscopy for characterizing mitochondrial morphology. A major challenge is the unbiased, accurate and quantitative assessment of complex mitochondrial morphology. The wide diversity of mitochondrial morphologies (elongated, fragmented, tubular, or mixed forms) complicates both quantitative evaluation and automated analysis (Collins et al., 2002). Autofluorescence, variable signal-to-noise, and background fluorescence can further confound automated detection. Manual segmentation can address some limitations but is subjective, low throughput, and time-consuming, requiring significant expertise. The advent of deep learning, particularly convolutional neural networks (CNNs) (Gonzalez, 2007), has enabled valuable advances in mitochondrial segmentation and quantification, but many existing tools remain limited in throughput, scalability, or robustness across diverse datasets and imaging conditions.

In response to the growing need for high-throughput and quantitative assessment of mitochondrial morphology and dynamics, we here describe the development of a High-throughput Mitochondrial Imaging Platform (H-MIP), combined with a deep learning segmentation tool, Mitochondrial Segmentation Network (MitoSegNet) (Fischer et al., 2020). This enables the automated analysis of millions of mitochondria in cells. By leveraging artificial intelligence (AI) for accurate segmentation and morphological classification, H-MIP offers a powerful and unbiased method for quantifying mitochondrial shape, length, and network complexity across large datasets in various experimental conditions with high sensitivity. This approach provides a robust indicator of mitochondrial health based on their morphology.

We first validated H-MIP by quantifying mitochondrial morphology in cells treated with mitochondrial division inhibitor 1 (Mdivi-1) (Manczak et al., 2019), confirming that length-associated metrics increase upon division inhibition. Recent studies highlight the importance of mitochondrial structure and function in vascular health (Demer & Tintut, 2008; Lee et al., 2020; Wang et al., 2020; Yu et al., 2012; Zeng et al., 2022). Consistent with this, we previously found that mitochondrial morphology is adversely affected in vascular smooth muscle cells (VSMCs) exposed to conditions that induce VC, a pathological process driven by abnormal calcium and phosphate (Pi) deposition as hydroxyapatite (HA) in the vessel wall, leading to vascular stiffening (Durham et al., 2018; Ortega et al., 2024). We therefore used H-MIP to investigate the morphology of mitochondria in VSMCs incubated under conditions favouring VC, and confirmed that the mitochondria are elongated.

Overall, H-MIP represents a significant advance in the quantitative analysis of mitochondrial morphology, combining high-throughput imaging with deep learning–based analytics for automated, objective assessment of mitochondrial morphology. Our studies demonstrate its ability to quantify key morphological changes associated with disease, providing a robust pipeline for both mechanistic discovery and high-content drug screening. By enabling unbiased, scalable, and sensitive evaluation of organelle health, H-MIP holds great promise for accelerating progress in mitochondrial biology, testing the development of mitochondrial-targeted therapies, and ultimately informing treatment strategies for multiple diseases where mitochondrial dysfunction is central.

## Results

### High-throughput imaging of mitochondria and finetuned MitoSegNet model

Fluorescence microscopy is a powerful tool for visualizing organelles within cells. To extract further information from the images, segmentation approaches can be utilized to distinguish objects of interest from background, commonly using thresholding approaches that identify pixels above a defined intensity threshold. Standard thresholding approaches, such as Otsu’s method, are commonly used to segment organelles, including mitochondria, in fluorescence images (Smith et al., 1979). Otsu’s method automatically determines an optimal threshold that effectively separates the foreground and background based on the assumption of a bimodal histogram (two distinct pixel groups, such as a dark background and bright mitochondria). Such intensity-based thresholding, however, can be limited in performance, particularly with varying signal intensity, signal-to-background and signal-to-noise ratios in diverse images (Valente et al., 2017). These limitations may lead to the omission of small or fragmented mitochondria, or conversely, to the erroneous merging of adjacent mitochondrial structures, resulting in inaccurate segmentation.

In our previous work, we developed a deep learning-based mitochondrial segmentation tool, MitoSegNet (Fischer et al., 2020), a modified U-Net based on fully CNN operations (Ronneberger et al., 2015), and demonstrated that it outperformed conventional methods in identifying mitochondria in fixed HeLa cells (Fischer et al., 2020). In the present study, we fine-tuned the MitoSegNet model and extended its application to analyze mitochondrial morphology in live cells which were imaged using a high-content Opera Phenix spinning-disk confocal microscope. The fine-tuned, high throughput MitoSegNet model (MitoSegNetH) is better adapted to the mitochondrial features characteristic of this experimental system, including the specific cell type, microscope type, and imaging configuration. As with the original MitoSegNet model, however, MitoSegNetH can be retrained on any mitochondria imaging dataset. As a model system, vascular cells were cultured in multi-well plates (96 well-plates, ***Figure 1A***), and imaged using a high-content Opera Phenix spinning-disk confocal microscope (***Figure 1B***). The acquired fluorescence images were segmented using MitoSegNetH and subsequently analyzed (***Figure 1C***), enabling quantification over 6 million mitochondria in live cells across 30,000 cells at maximum capacity (***Figure 1D***). After segmentation with MitoSegNetH, each mitochondrial object was measured for 11 morphological features. These features represent a deliberately selected subset that capture complementary and biologically interpretable aspects of mitochondrial network organisation, while minimising redundancy. Among these, we focused primarily on 4 mean values of 1) major axis length, 2) total branch length per mitochondrion, 3) number of branches, and 4) perimeter to further characterize mitochondrial morphology and assess changes under different conditions (***Figure 1E***).

**Figure 1.**
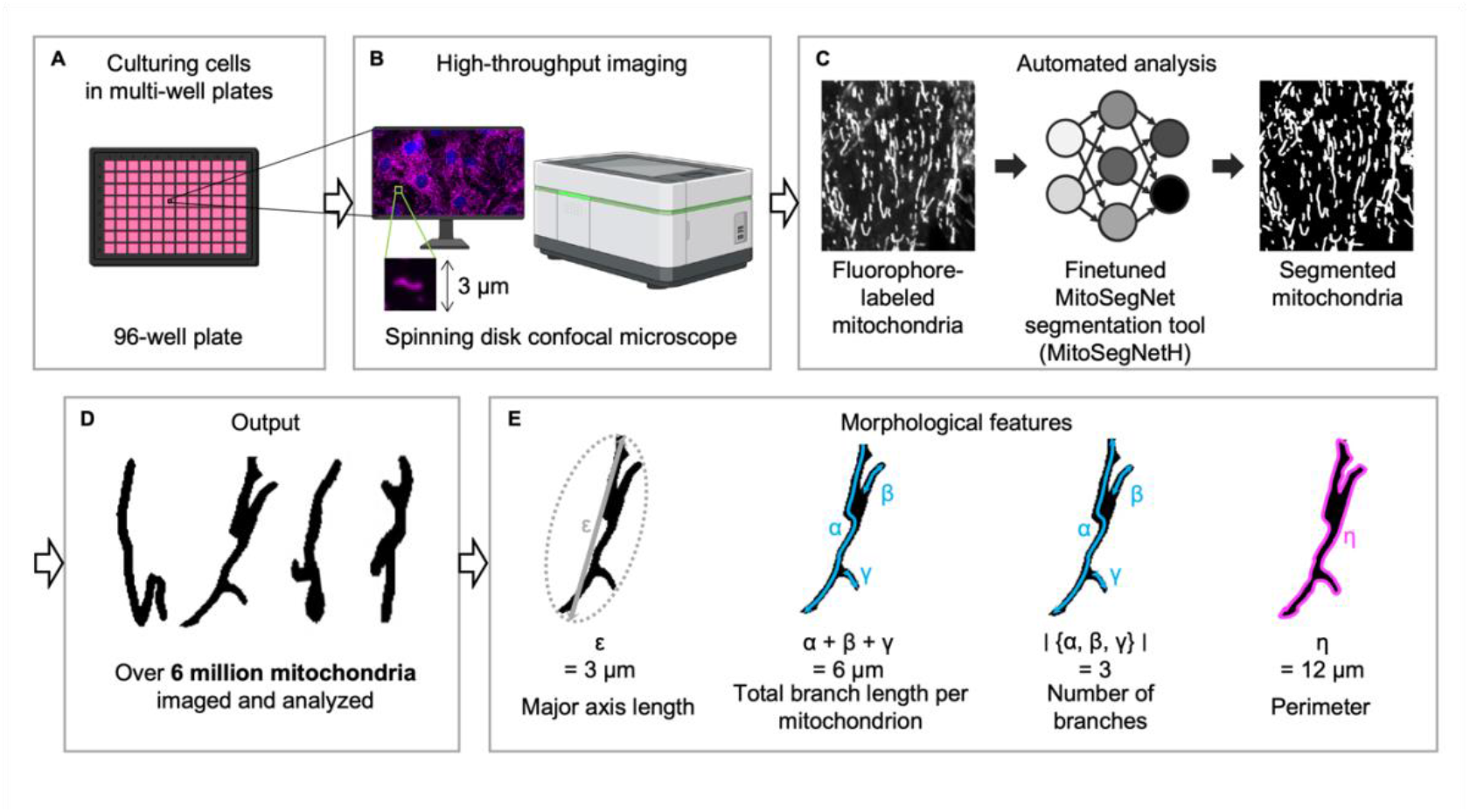
AI-assisted H-MIP enables high-throughput quantification of millions of individual mitochondrial features in live cells. Schematic H-MIP and AI pipeline. **A**. Cells were incubated on multi-well plates. **B**. Cells were imaged using a high-content Opera Phenix spinning-disk confocal microscope. **C**. The acquired images were subsequently segmented using the fine-tuned MitoSegNet model (MitoSegNetH) and their morphology analyzed. **D**. At maximum capacity, over 6 million mitochondria in live VSMCs can be analyzed. **E**. Representative mitochondrial features extracted from the MitoSegNetH package.

This workflow allowed us to quantify millions of mitochondrial structures and extract key morphological descriptors in a high-throughput manner, establishing MitoSegNetH as a powerful tool for mitochondrial segmentation and ultimately as a robust platform for assessing mitochondrial morphological changes.

### Benchmarking MitoSegNetH with H-MIP

To benchmark the MitoSegNetH (see Methods for details), fluorescence images with varying mitochondria densities (sparse: 0.13 ± 0.03 mitochondrial area/μm^2^, medium: 0.21 ± 0.05 mitochondrial area/μm^2^, and dense: 0.34 ± 0.11 mitochondrial area/μm^2^) were segmented using previously reported open-source algorithms: Otsu-based thresholding, Nellie (Lefebvre et al., 2025), MoDL (Ding et al., 2025), and MitoSegNet, as well as the MitoSegNetH (dense condition in ***Supplementary Figure 1*** shows all densities.). When compared with the ground truth human-annotated image, MitoSegNetH successfully segmented and distinguished intricate mitochondria across all density levels, including under the condition of dense, highly interconnected networks, by accurately identifying mitochondrial junctions and endpoints (***Figure 1A*** and ***Supplementary Figure 1***). MitoSegNetH achieved the highest performance in terms of accuracy and Dice coefficient combined with showing the smallest standard deviation, significantly outperforming Otsu-based thresholding, Nelli, MoDL, and the previous MitoSegNet algorithm. Each performance value at the different mitochondrial densities is also shown in ***Supplementary Table 1*** and ***Supplementary Figure 1***. MitoSegNetH provides a substantial improvement over conventional thresholding approaches, enabling accurate and automated segmentation as well as comprehensive morphological analysis of mitochondria in high-content fluorescence microscopy datasets. Also, MitoSegNetH markedly outperforms existing open-source mitochondrial segmentation methods across all tested density conditions. It delivers the highest accuracy and Dice coefficient scores with the lowest variability, and uniquely maintains reliable performance even in dense and highly interconnected networks. Together, these results establish MitoSegNetH as a robust and precise tool for quantitative segmentation of complex mitochondrial architectures in VSMCs.

### Assessment of changes in mitochondrial morphology

To demonstrate the ability of H-MIP to measure changes in mitochondrial morphology, we treated VSMCs with increasing concentrations of the mitochondrial division inhibitor 1 (Mdivi-1) (***Figure 3A***), a small molecule which inhibits mitochondrial fission mediated by dynamin-related protein 1 (Drp1) (***Figure 3B***) (Rosdah et al., 2022). Consistent with earlier work demonstrating that either genetic deletion or pharmacological inhibition of Drp1 promotes mitochondrial elongation (Rosdah et al., 2022; Tang et al., 2024), we anticipated a dose-dependent increase in mitochondrial length upon Mdivi-1 treatment.

Quantitative high-throughput imaging and analysis revealed significant increases in the major axis length following 100 µM Mdivi-1 treatment (***Figure 3C***). Additional morphological parameters, such as the total branch length per mitochondrion (***Figure 3D***), number of branches (***Figure 3E***), and perimeter (***Figure 3F***) also increased before plateauing at 50-75 µM, indicating a saturating structural response at higher concentrations. These observations align with the established morphological signature of Drp1 inhibition (Tang et al., 2024) and confirm H-MIP’s ability to capture gradations in mitochondrial network elongation. Furthermore, our findings are consistent with recent studies reporting that Mdivi-1 induces pronounced mitochondrial elongation and network hyperfusion in diverse cell types, including fibroblasts, cardiomyocytes, and neuronal cells (Hoque et al., 2018; Lin et al., 2024; Ruiz et al., 2018).

**Figure 2.**
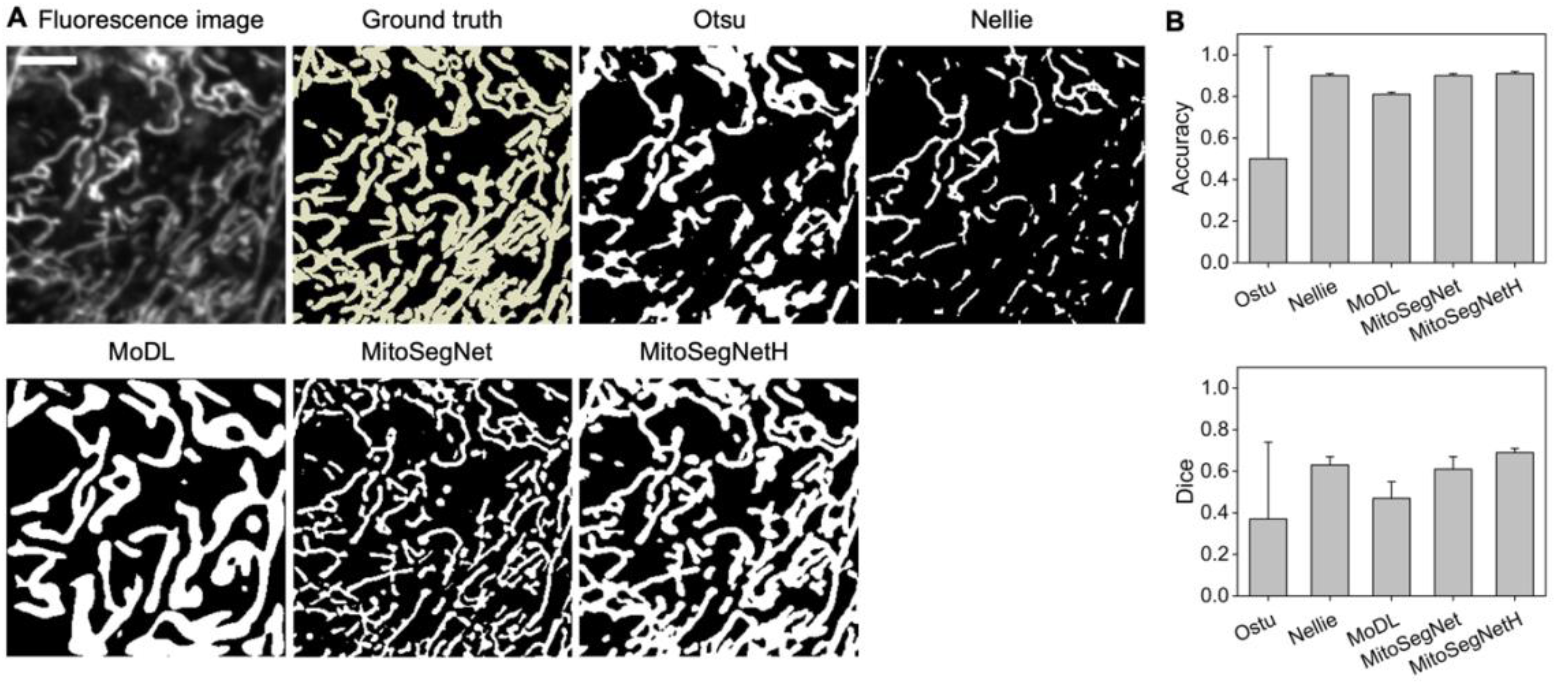
MitoSegNetH outperforms other mitochondria segmentation methods. **A**. A comparison of segmented mitochondria from Otsu thresholding, Nellie (Lefebvre et al., 2025), MoDL (Ding et al., 2025), MitoSegNet (Fischer et al., 2020) and MitoSegNetH with the ground truth. Scale bar = 5 μm. **B**. Comparison of segmentation performance in dense condition. n = 6 images from different density levels of mitochondria (see **Supplementary Figure 1A** and **Supplementary Table 1**).

**Figure 3.**
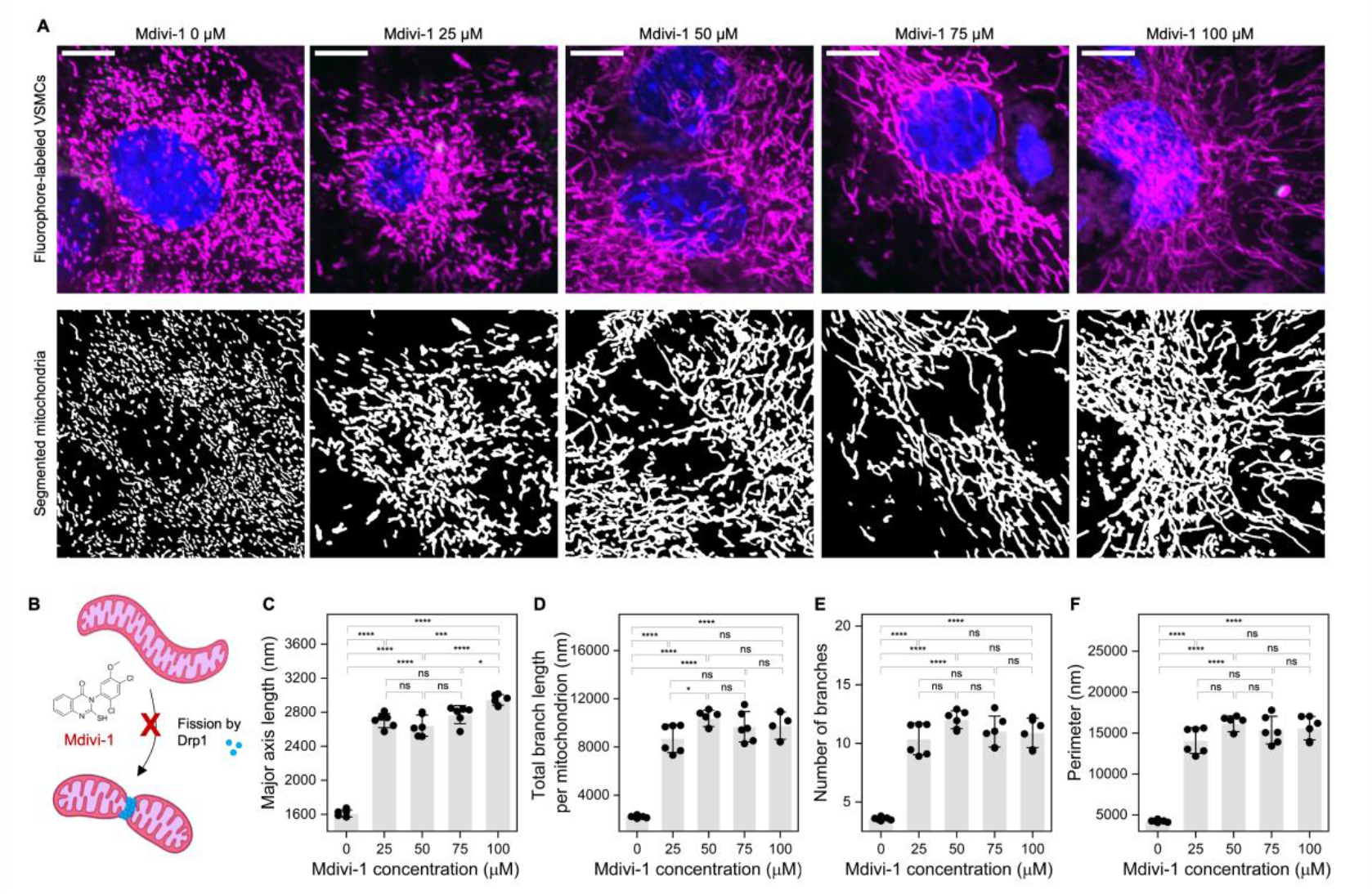
H-MIP demonstrates that the mitochondrial division inhibitor 1 (Mdivi-1) modulates mitochondrial morphology by promoting elongation. **A**. Representative images of fluorophore-labeled VSMCs and segmented mitochondria treated with increasing concentrations of Mdivi-1. Mitochondria (MitoLite, magenta) and nuclei (Hoechst, blue). Scale bar 10 μm. **B**. Mdivi-1 inhibits the key fission protein dynamin-related protein 1 (Drp1). **C-F**. Morphology changes significantly with an increasing concentration of Mdivi-1 from 0 to 100 μM. n = 312 images per condition. Error bars indicate standard deviation (SD) from 6 biological repeats, and statistical significance was determined using one-way analysis of variance (ANOVA) with post hoc Tukey pairwise means comparisons, ns, not significant, * P < 0.05, ** P < 0.01, *** P < 0.001, **** P < 0.0001. **C**. Major axis length increased from 1,610 ± 41 nm to 2,944 ± 62 nm, **D**. Total branch length per mitochondrion increased from 2,206 ± 105 nm to 10,362 ± 676 nm and plateaued at 50 - 75 μM, **E**. The number of branches and **F**. perimeter increased from 3.6 ± 0.2 to 12.0 ± 0.8 and from 4,249 ± 149 nm to 16,144 ± 975 nm, respectively, and plateaued at 50 - 75 μM.

A further advantage of H-MIP with MitoSegNetH is the ability to analyze entire populations of mitochondria, and this revealed considerable heterogeneity in mitochondrial architecture (***Supplementary Figure 2***). In particular, higher Mdivi-1 concentrations were associated with increased heterogeneity, indicating that individual mitochondria occupy a broad range of dynamic states. Importantly, the increased population of mitochondria with longer major axis lengths implies a shift in dynamic equilibrium, in which fusion events dominate and fragmentation no longer efficiently counterbalances elongation. Collectively, these distributions reflect altered fusion-fission dynamics rather than uniform morphological changes.

Taken together, the observed mitochondrial elongation and progressive changes across multiple quantitative morphological features indicate that Mdivi-1 promotes a more elongated and interconnected mitochondrial architecture by Drp1 inhibition. Our findings highlight the sensitivity and robustness of H-MIP for high-throughput detection of subtle, biologically relevant structural alterations across diverse experimental conditions.

### Assessment of mitochondrial morphology in an established *in vitro* model of vascular calcification

Mitochondrial dysfunction is a critical driver of VC (Phadwal et al., 2021; Wang et al., 2020; Yu et al., 2012), with alterations in mitochondrial morphology closely associated with mitochondrial dysfunction. To evaluate the mitochondrial morphology associated with VC, we combined our previously established calcification model (Zhu et al., 2014, 2016) with H-MIP. In this model, VSMCs were exposed to high inorganic phosphate (3 mM Pi) to mimic the VC environment and were cultured in 96-well plates for up to 14 days (***Figure 4A***). Calcification was quantified using Alizarin Red staining (***Figure 4B***), and the *o*-Cresolphthalein complexone assay (***Figure 4C***), whereby the *o*-Cresolphthalein complexone reacts with free calcium under alkaline conditions to produce a purple color, which can be measured spectrophotometrically. Both methods revealed a significant increase in calcium deposition in VSMCs cultured under high Pi conditions compared to control cells.

**Figure 4.**
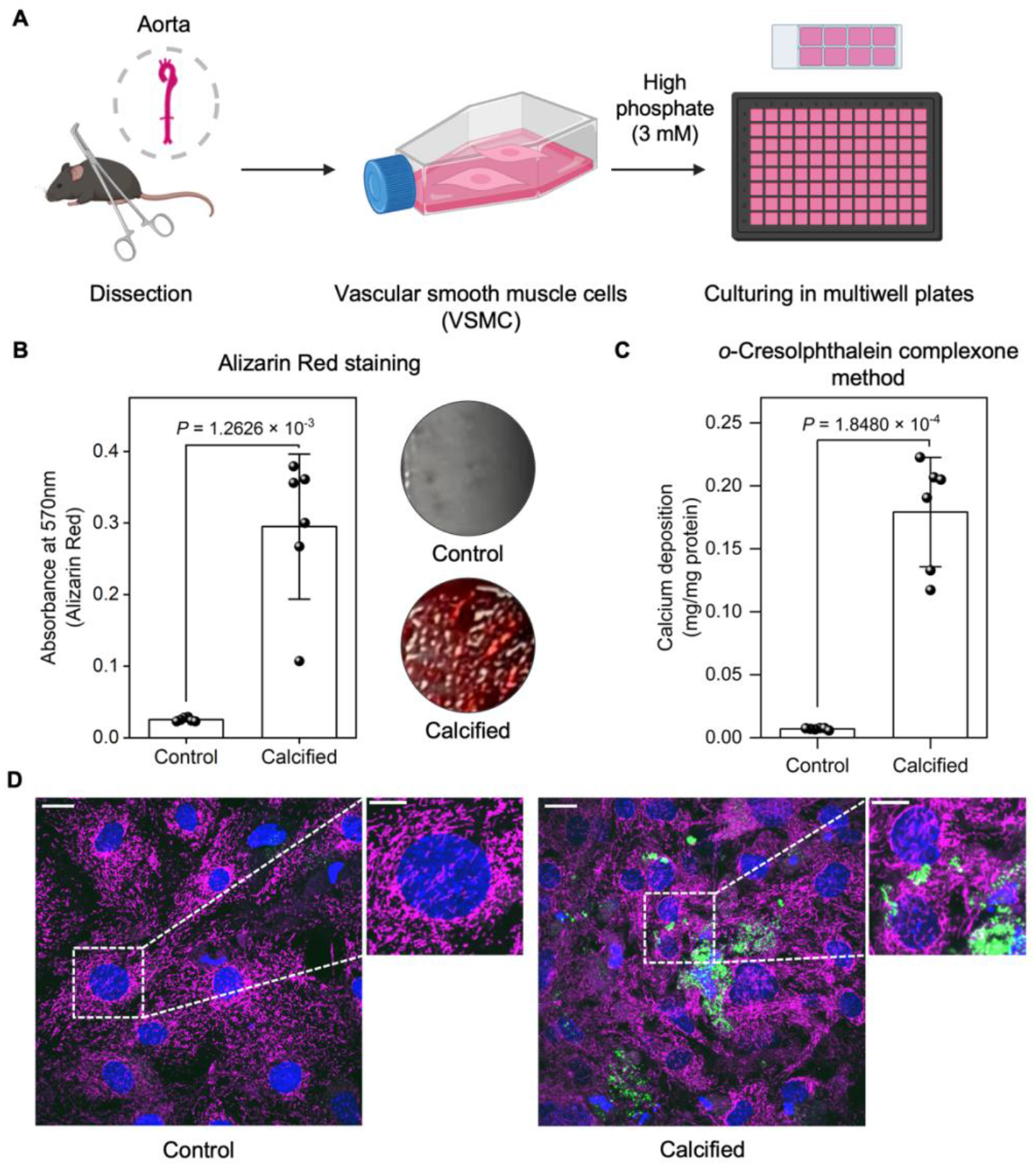
Use of H-MIP to investigate mitochondrial morphology associated with VC as an exemplar disease model. **A**. Primary VSMCs were isolated from aortae of male C57B/6 mice. VSMCs were first cultured in T25 and T75 flasks and then were seeded at a density of 1.0 × 10^4^/well on 96-well plates. **B**. Confirmation of calcium deposition in calcifying VSMCs using Alizarin Red staining on 96-well plates. Control VSMCs (grey), calcifying VSMCs (red), red color is an indicator of calcium deposition in calcifying VSMCs. Error bars indicate SD of 6 biological repeats, and statistical significance was determined using an unpaired t-test, P = 1.2626 × 10^-3^. **C**. Quantification of calcium deposition in control and calcifying VSMCs using o-Cresolphthalein complexone method performed on 96 well-plates. A significantly higher calcium level was observed in calcifying VSMCs (0.1792 ± 0.0434) compared to the value in control VSMCs (0.0070 ± 0.0008). Error bars indicate SD of 6 biological repeats, and statistical significance was determined using t-test, P = 1.8480 × 10^-4^. **D**. Representative images of control and calcifying VSMCs using spinning disk confocal microscopy. A solution of fluorescein-bisphosphonate (2 μM) was added for staining calcium deposition (green). MitoLite was diluted 1:500 and then added for staining mitochondria (magenta). Hoechst was used for staining nucleus (blue). Scale bar 20 μm. Inset scale bar: 10 μm.

To further assess calcification, we visualized the deposition of HA, the primary mineral component that forms in VC, using fluorescein-bisphosphonate, a novel HA-specific fluorescent probe developed by our laboratory (Sim et al., 2018) (***Figure 4D***). Prominent HA deposition was observed in calcifying VSMCs (green color), while control cells exhibited negligible signal. These results validate the robustness of our Pi-induced VC model in replicating the VC environment and provide a reliable platform for investigating mitochondrial health through morphological change during calcification.

Having validated the calcification model, we next sought to evaluate mitochondrial morphological changes associated with VSMC calcification using H-MIP. To achieve this, we incubated VSMCs with increasing concentrations of Pi (1.5, 2, 2.5, and 3 mM). This led to a progressive accumulation of HA, as evidenced by representative images (***Figure 5A***) and quantification (***Figure 5B***). In our previous work, we reported mitochondrial elongation in fixed VSMCs treated with 3 mM Pi compared to control VSMCs (Phadwal et al., 2023). Consistent with these findings, the present study revealed a progressive elongation of mitochondrial morphology in live calcifying VSMCs cultured with increasing Pi concentrations from 1.5 mM to 3 mM. Significant increases in metrics associated with morphological elongation were detected in VSMCs treated with 3 mM Pi compared to those treated with 1.5 mM Pi (***Figure 5C-F***).

**Figure 5.**
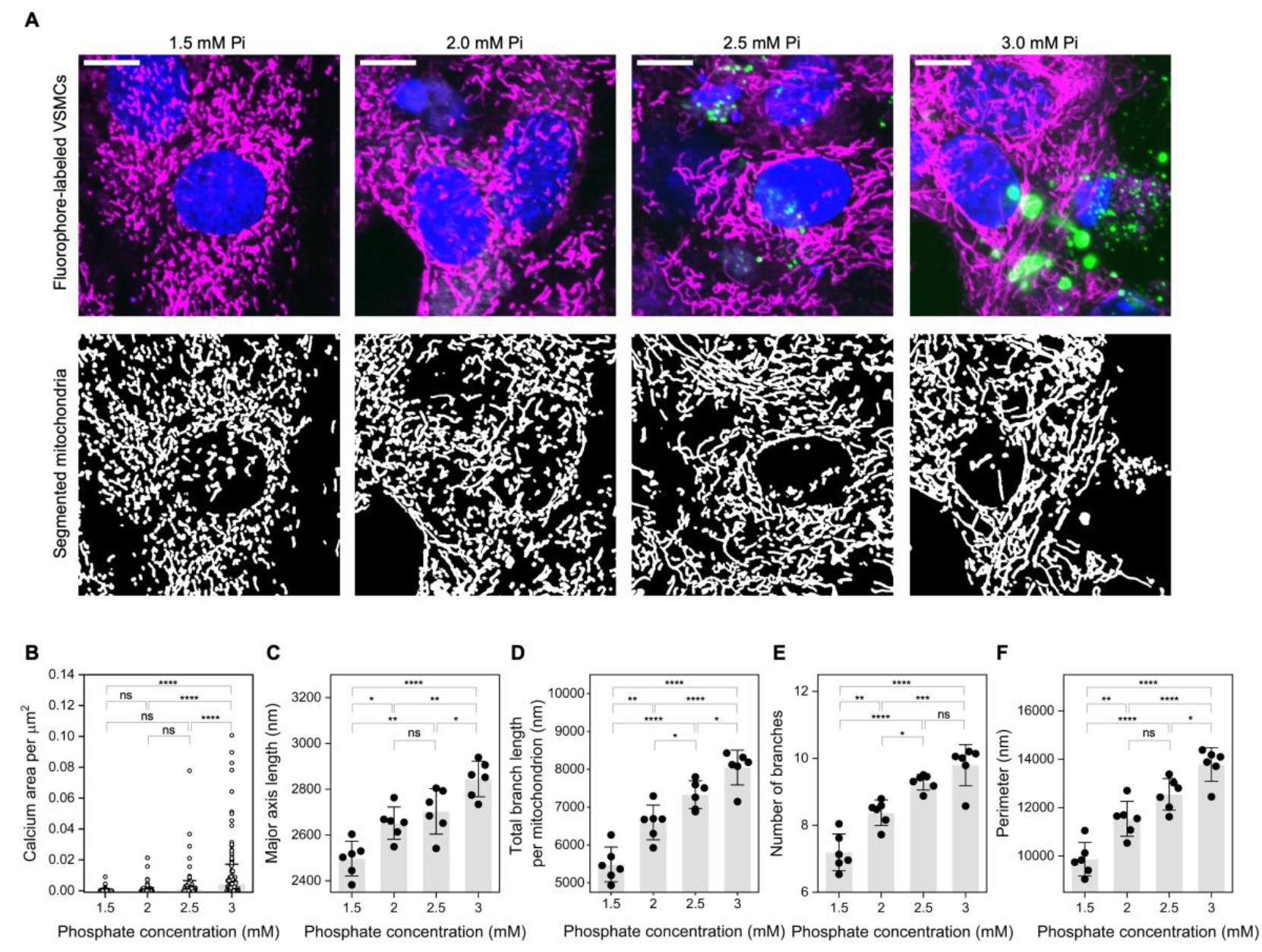
H-MIP captures progressive elongation of mitochondrial morphology in live VSMCs cultured in calcification media with elevated Pi. **A**. Representative images of fluorophore-labeled VSMCs and AI segmented mitochondria cultured with different Pi concentrations. Mitochondria (magenta), nucleus (blue), and calcium deposition HA (green). Scale bar 10 μm. n = 300 images per condition. **B**. Quantification of calcium deposition in calcifying VSMCs in fluorescence images. SD values represent independent measurements from separate fields of view (n = 300). Statistical significance was determined by ANOVA with post hoc Tukey pairwise means comparisons, ns, not significant, * P < 0.05, ** P < 0.01, *** P < 0.001, **** P < 0.0001. **C**. Major axis length, **D**. Total branch length per mitochondrion **E**. The number of branches. **F**. Perimeter. Error bars indicate SD of 6 biological repeats, and statistical significance was determined using ANOVA with post hoc Tukey pairwise means comparisons, ns, not significant, P > 0.05, * P < 0.05, ** P < 0.01, *** P < 0.001, **** P < 0.0001.

Notably, VSMCs exposed to the highest concentration of Pi (3 mM) showed a higher proportion of elongated mitochondria compared with cells cultured under lower Pi conditions, as shown by histograms of major axis length for individual mitochondria (***Supplementary Figure 3A***). To further characterize mitochondrial length distributions, we constructed cumulative distribution functions, revealing a Pi concentration-dependent rightward shift most pronounced at 3 mM Pi. (***Supplementary Figure 3B***). These findings indicate an increased proportion of longer mitochondria under higher calcifying conditions and are consistent with progressive elongation and expansion of the mitochondrial network in response to elevated Pi. Together, these results indicate that Pi-induced calcification drives distinct alterations in mitochondrial morphology, reflecting an adaptive remodeling characterized by elongation, increased branching and network expansion, which may represent an early morphological hallmark of mitochondrial dysfunction during VC.

## Discussion

Mitochondrial structure is not merely a downstream consequence of altered function but can correlate with - and in some contexts actively drive - functional changes in cellular bioenergetics and signaling (Benard et al., 2008). Alterations in mitochondrial architecture, particularly at the level of cristae organization and network connectivity, can impose rate-determining constraints on oxidative phosphorylation and metabolic flexibility, and are increasingly implicated across a wide spectrum of diseases, including VC, PD, cancer, and CKD (Huang et al., 2024; Klemmensen et al., 2024; Moradi Vastegani et al., 2023). Despite this, systematic interrogation of mitochondrial structural remodeling has been limited by the lack of scalable, quantitative, and unbiased analytical approaches.

To address this gap, we developed H-MIP, a high-throughput platform that enables visualization and quantitative assessment of mitochondrial morphology at single-mitochondrion resolution, providing a sensitive structural readout of mitochondrial health. By integrating high-sensitivity spinning-disk confocal microscopy with the AI-driven MitoSegNetH analysis pipeline, H-MIP enables large-scale, automated quantification of mitochondrial morphology across diverse experimental conditions. This approach dramatically extends imaging and analytical capacity, permitting interrogation of up to ∼800,000 cells and ∼150 million mitochondria in fixed samples at maximum throughput, representing an approximately two-order-of-magnitude improvement over conventional approaches. Benchmarking against Otsu’s thresholding and recently developed open-source algorithms, including Nelli and MoDL, demonstrated superior accuracy and robustness of H-MIP across complex mitochondrial phenotypes. Using a panel of 11 quantitative morphological features encompassing size, length, and network connectivity H-MIP enables comprehensive, multidimensional profiling of mitochondrial structure. To our knowledge, this study represents the first demonstration of high-throughput imaging combined with AI-based analysis of mitochondrial morphology at this scale.

The sensitivity and robustness of the platform were validated using pharmacological modulation of mitochondrial dynamics. Treatment of healthy VSMCs with increasing concentrations of the mitochondrial fission inhibitor Mdivi-1 resulted in dose-dependent mitochondrial elongation, consistent with inhibition of Drp1-mediated fission (Rosdah et al., 2022; Tang et al., 2024; Z. Wang et al., 2024). Importantly, H-MIP captured subtle and progressive morphological shifts across large cell populations, highlighting its suitability for high-content screening applications and quantitative assessment of dynamic mitochondrial responses to pharmacological perturbation.

We further applied H-MIP to a phosphate-induced VC model, revealing progressive mitochondrial elongation and hyperfusion in VSMCs treated with increasing Pi concentrations. These findings are consistent with previous reports from our laboratory using low-throughput, non-automated approaches (Phadwal et al., 2023), and align with recent studies linking mitochondrial hyperfusion in calcifying VSMCs to elevated reactive oxygen species, loss of membrane potential, and impaired mitophagy (Phadwal et al., 2021; Zeng et al., 2022). Together, these observations suggest that elongated mitochondrial morphology serves as a structural signature of mitochondrial stress and dysfunction during calcification.

Beyond disease modeling, the ability to quantify mitochondrial morphology at scale has important implications. By resolving mitochondrion-to-mitochondrion variability within single cells, H-MIP provides insights into subcellular heterogeneity that are inaccessible to bulk biochemical assays, allowing structural phenotypes to be linked to functional outcomes. This approach enables systematic investigation of how mitochondrial shape encodes metabolic state, stress adaptation, and cell fate decisions, advancing our understanding of structure– function relationships in mitochondrial biology. Furthermore, high-throughput morphological profiling enables unbiased screening of potential therapeutics for their effects on mitochondrial dynamics, not only facilitating identification of compounds that restore healthy network architecture but also identifying crucial off-target effects on mitochondrial function.

In summary, H-MIP represents a scalable and sensitive platform for high-content analysis of mitochondrial morphology, with broad applicability to drug screening, disease modeling, and mechanistic studies. By combining AI-driven segmentation with high-throughput imaging at single-mitochondrion resolution, H-MIP captures structural heterogeneity as a biologically meaningful dimension of mitochondrial health, thereby supporting precision medicine and therapeutic strategies targeting mitochondrial dysfunction.

## Methods

### Preparation of VSMCs

Primary VSMCs were prepared from 6-week-old wild-type (WT) C57B/6 mice and cultured in growth medium as previously described (Zhu et al., 2016). Briefly, the adventitia was carefully removed, and the aorta was opened to reveal the endothelial layer using a dissection microscope. Tissues from eight mice were pooled and incubated with 1 mg/mL trypsin (Invitrogen, Paisley, UK) for 10 minutes to remove any remaining adventitia and endothelium. The tissues were then incubated overnight at 37°C in a humidified atmosphere of 5% CO_2_ in growth medium consisting of containing Minimum Essential Medium Eagle alpha modification (α-MEM; Life Technologies) supplemented with 10% FBS and 1% gentamicin (all from Invitrogen). Following this, the tissues were digested with 425 U/ml collagenase type II for 5 hours. The resulting cell suspensions were centrifuged at 2000 x g for minutes, and the cell pellet was washed and resuspended in the same growth medium. Isolated VSMCs were cultured in growth medium for two passages in T25 tissue culture flasks (Greiner Bio-one, GmbH, Frickenhausen, Baden-Wurttemberg, Germany) coated with 0.25 μg/cm^2^ laminin (Sigma, Poole, UK) to support the maintenance of the contractile differentiation state.

### VSMC Cell Culture

VSMCs were maintained in growth medium of α-MEM supplemented with 10% FBS (Invitrogen) and 1% gentamicin (Invitrogen) and cultured at 37°C in a humidified 5% CO_2_ Incubator. For experiments, cells were first seeded into 96-well plates and allowed to grow to 70-80% confluency before beginning respective treatments.

### Induction of VSMC calcification

Calcification was induced as previously described (Phadwal et al., 2023; Zhu et al., 2016). In brief, for VSMCs, cells were grown to confluence (day 0) and switched to calcification medium, which was prepared by adding 1M inorganic Pi (a mixture of NaH_2_PO_4_ and Na_2_HPO_4_, pH 7.4) (Sigma-Aldrich), to reach a final concentration of 3 mM Pi in α-MEM. Cells were incubated for up to 14 days in 5% CO_2_ at 37°C and medium was changed every second/third day.

### Determination of VSMC calcification

Calcium content was quantified by HCl leaching and the *o*-Cresolphthalein complexone method, as described previously (Sim et al., 2018). Briefly, cells were rinsed twice with phosphate buffered saline (PBS) and incubated with 0.6 M HCl at room temperature for 24 hours. Free calcium was determined colormetrically by a stable interaction with phenolsulphonethalein using a commercially available kit (Randox Laboratories Ltd., County Antrim, UK), corrected for total protein concentration (Bio-Rad Laboratories Ltd, Hemel Hempstead, UK), and presented as a fold change compared with control. Calcium deposition was evaluated by Alizarin Red staining as previously described (Sim et al., 2018). Cells were washed twice with PBS, fixed in 4% paraformaldehyde for 5 minutes at 4°C, stained with 2% Alizarin Red (pH 4.2) for 5 minutes at room temperature and rinsed with distilled water.

### Preparation of live VSMCs and cell imaging with Opera Phenix

VSMCs were seeded at a density of 1.0 × 10^4^/well in 96-well ibidi plates. For Mdivi-1 treatment, the cells were exposed to different concentration of Mdivi-1 upon reaching confluency then cultured for up to 14 days and and medium was changed every second/third day. For calcification experiments, cells were induced to calcify upon reaching confluency and cultured for up to 14 days. At day 14, 2 μM of fluorescein-bisphosphonate was added and incubated in 5% CO_2_ at 37°C for 1 hour. Cells were then washed three times with PBS, MitoLite™ Red FX600 (AAT Bioquest) was diluted 1:500 and then added. The cells were incubated in 5% CO_2_ at 37°C for 30 minutes. Cells were then washed three times with PBS, and cells were stained with Hoechst (H3570, Thermo Scientific) for 5 minutes and then washed three times with PBS before they were ready for imaging. After staining the VSMCs, the VSMCs in multi-well plate were imaged using a high-throughput imaging system, Opera Phenix confocal microscope (Perkin Elmer). For each condition, up to 312 images were taken.

### Mitochondria image segmentation

Fluorescence images of mitochondria captured using Opera Phenix spinning-disk confocal microscope were used as input for the analysis. MitoSegNet (Fischer et al., 2020) is a U-Net (Ronneberger et al., 2015) based deep learning tool to segment fluorescently labeled mitochondria images. Using MitoSegNet model (Fischer et al., 2020) on data that are different from those it was trained on, generates sub-optimal results with very thin boundaries and missed structures. Therefore, MitoSegNet was fine-tuned using our experimental image data to create the MitoSegNetH model. Fine-tuning was performed using image-label pairs that were manually annotated with Labelme (Russell et al., 2008). MitoSegNetH was MitoSegNet fine-tuned using an expert-annotated VSMC image ROI of size 400 × 400 pixels. By applying data augmentation techniques, we obtained a training set of 128 training images and 32 validation images of size 256 × 256 (the input size of MitoSegNet). To reduce the presence of fragmented mitochondria across the axial dimension, training images were maximum intensity projections of 3 slices from z-stacks of images section.

### Morphological analysis

The segmented images of mitochondria were processed to extract morphological features of individual mitochondria using the scipy (Van Der Walt et al., 2014) and skan (Sundar et al., 2003) libraries. Area, perimeter, solidity, major axis length, minor axis length, and eccentricity were measured using the region labelling function from scipy, and branch length, branch length per mitochondrion and curvature index were measured using the skeleton analysis functions from the skan library.

## Acknowledgements

VEM is a member of the International Network on Ectopic Calcification (INTEC **· itnintec.com**). For the purpose of open access, the author has applied a CC-BY public copyright license to any Author Accepted Manuscript version arising from this submission.

