## Supplemental information for "Live high-content imaging with automated analysis reveals mitochondrial changes during vascular calcification"

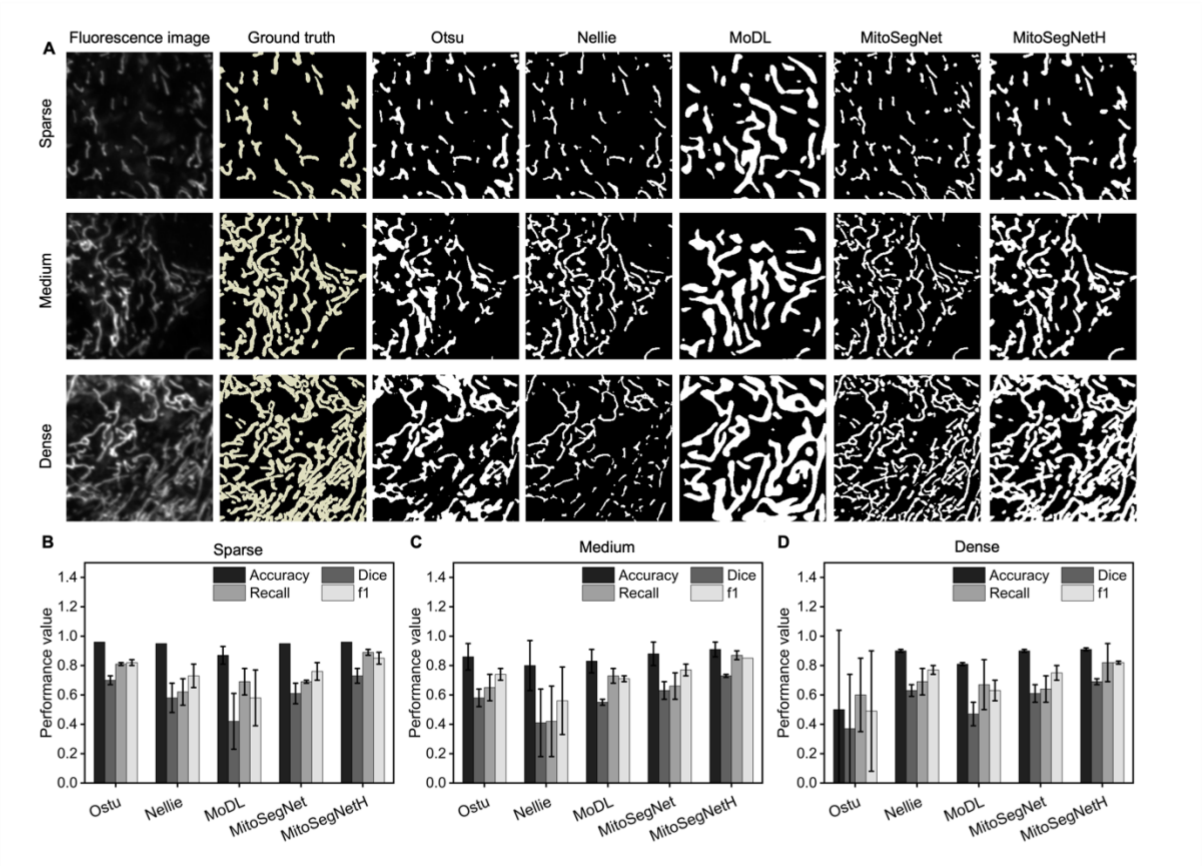


**Supplementary Figure 1. A. Fine-tuned MitoSegNetH outperformed the alternative reported algorithms i.e., Otsu, Nellie, MoDL, and MitoSegNet when analyzing varying mitochondrial densities.** When compared with the ground truth, only MitoSegNetH model successfully segmented and distinguished intricate mitochondria across all density levels, including dense, highly interconnected networks, by accurately identifying mitochondrial junctions and endpoints. **B-D** Segmentation performance comparison of MitoSegNetH, open-source deep learning algorithms that are retrained with confocal image dataset, i.e., Otsu. Nellie, MoDL, MitoSegNet and MitoSegNetH. n = 6 images representing varying mitochondrial densities. The fine-tuned MitoSegNetH achieved the highest values in most of performances, overall outperforming Ostu-based thresholding Nelli, MoDL, and MitoSegNet. The exact values are shown in the Supplementary Table 1.

**Supplementary Table 1.** Comparison of segmentation performance among Otsu thresholding, Nellie (Lefebvre et al., 2025), MoDL (Ding et al., 2025), MitoSegNet (Fischer et al., 2020) and MitoSegNetH.

| | Method | Accuracy | Dice | Recall | f1 |
|---|---|---|---|---|---|
| **Sparse** | **Ostu** | 0.96 ± 0.00 | 0.70 ± 0.03 | 0.81 ± 0.01 | 0.82 ± 0.02 |
| | **Nellie** | 0.95 ± 0.00 | 0.58 ± 0.10 | 0.62 ± 0.09 | 0.73 ± 0.08 |
| | **MoDL** | 0.87 ± 0.06 | 0.42 ± 0.19 | 0.69 ± 0.09 | 0.58 ± 0.19 |
| | **MitoSegNet** | 0.95 ± 0.00 | 0.61 ± 0.07 | 0.69 ± 0.01 | 0.76 ± 0.06 |
| | **MitoSegNetH** | 0.96 ± 0.00 | 0.73 ± 0.05 | 0.89 ± 0.02 | 0.85 ± 0.04 |
| **Medium** | **Ostu** | 0.86 ± 0.09 | 0.58 ± 0.06 | 0.65 ± 0.09 | 0.74 ± 0.04 |
| | **Nellie** | 0.80 ± 0.17 | 0.41 ± 0.23 | 0.42 ± 0.24 | 0.56 ± 0.23 |
| | **MoDL** | 0.83 ± 0.08 | 0.55 ± 0.02 | 0.73 ± 0.05 | 0.71 ± 0.02 |
| | **MitoSegNet** | 0.88 ± 0.08 | 0.63 ± 0.06 | 0.66 ± 0.09 | 0.77 ± 0.04 |
| | **MitoSegNetH** | 0.91 ± 0.05 | 0.73 ± 0.01 | 0.87 ± 0.03 | 0.85 ± 0.00 |
| **Dense** | **Ostu** | 0.50 ± 0.54 | 0.37 ± 0.37 | 0.60 ± 0.25 | 0.49 ± 0.41 |
| | **Nellie** | 0.90 ± 0.01 | 0.63 ± 0.04 | 0.69 ± 0.09 | 0.77 ± 0.03 |
| | **MoDL** | 0.81 ± 0.01 | 0.47 ± 0.08 | 0.67 ± 0.17 | 0.63 ± 0.07 |
| | **MitoSegNet** | 0.90 ± 0.01 | 0.61 ± 0.06 | 0.64 ± 0.09 | 0.75 ± 0.05 |
| | **MitoSegNetH** | 0.91 ± 0.01 | 0.69 ± 0.02 | 0.82 ± 0.13 | 0.82 ± 0.01 |

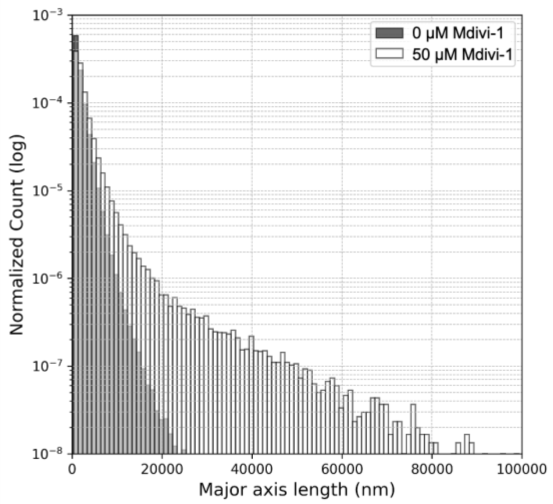


**Supplementary Figure 2. AI assisted H-MIP detects the heterogeneous morphological features of mitochondria.** Comparison of major axis length of individual mitochondrial morphology between 0 μM Mdivi-1 and 50 μM Mdivi-1 treatment in histograms.

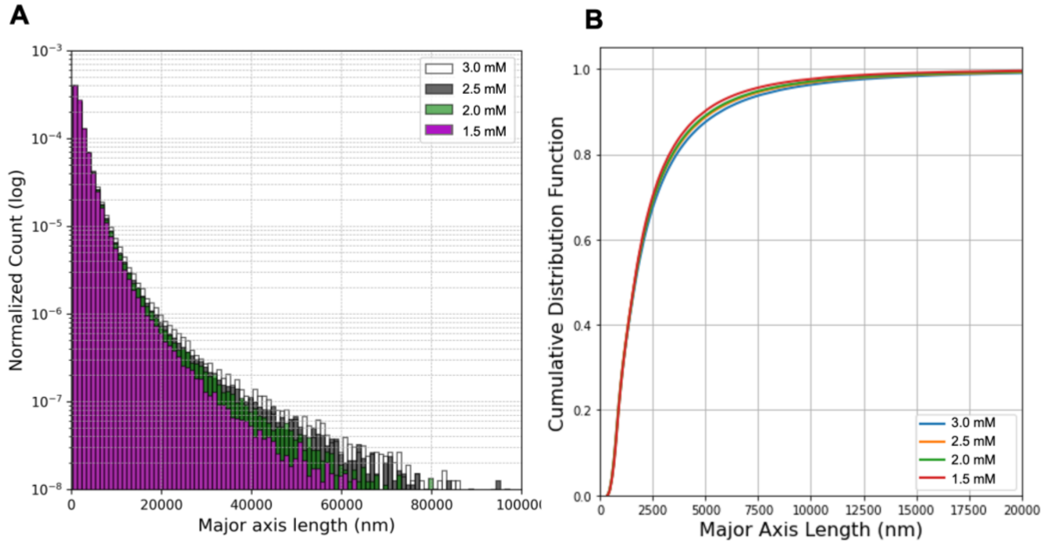


**Supplementary Figure S3. AI assisted H-MIP detects the heterogeneous morphological features of mitochondria.** Comparison of major axis length of individual mitochondrial morphology with elevated Pi in histograms (A) and cumulative distribution frequency (B).
